# Pan-genome-scale network reconstruction: a framework to increase the quantity and quality of metabolic network reconstructions throughout the tree of life

**DOI:** 10.1101/412593

**Authors:** Kevin Correia, Radhakrishnan Mahadevan

## Abstract

A genome-scale network reconstruction (GENRE) represents the knowledgebase of an organism and can be used in a variety of applications. The drop in genome sequencing costs has led to an increase in sequenced genomes, but the number of curated GENRE’ s has not kept pace. This gap hinders our ability to study physiology across the tree of life. Furthermore, our analysis of yeast GENRE’ s has found they contain significant commission and omission errors, especially in central metabolism. To address these quantity and quality issues for GENRE’ s, we propose open and transparent curation of the pan-genome, pan-reactome, pan-metabolome, and pan-phenome for taxons by research communities, rather than for a single species. We outline our approach with a Fungi pan-GENRE by integrating AYbRAH, our ortholog database, and AYbRAHAM, our new fungal reaction database. This pan-GENRE was used to compile 33 yeast/fungi GENRE’ s in the Dikarya subkingdom, spanning 600 million years. The fungal pan-GENRE contains 1547 orthologs, 2726 reactions, 2226 metabolites, and 10 compartments. The strain GENRE’ s have a wider genomic and metabolic than previous yeast and fungi GENRE’ s. Metabolic simulations show the amino acid yields from glucose differs between yeast lineages, indicating metabolic networks have evolved in yeasts. Curating ortholog and reaction databases for a taxon can be used to increase the quantity and quality of strain GENRE’ s. This pan-GENRE framework provides the ability to scale high-quality GENRE’ s to more branches in the tree of life.

## 1 INTRODUCTION

A genome-scale network reconstruction (GENRE) represents the knowledgebase of an organism and has a variety of applications. Studies have used GENRE’ s in metabolic simulations to uncover new enzyme-metabolite interactions (Hackett et al., 2016) and understand the pharmacokinetic response of drugs in the human body (Thiele et al., 2017), but they also can be used as a scaffold for omics integration (Österlund et al., 2013).

Current practices limit the quantity and quality of GENRE’ s, and their application to study metabolism throughout the tree of life. First, the drop in genome sequencing costs and the dominant paradigm of one research team for every GENRE has led to a growing gap between genome sequences and curated GENRE’ s. Second, most GENRE efforts are directed to a handful of organisms with existing GENRE’ s (Monk et al., 2014); the yeast/fungal GENRE community primarily focuses on industrially-relevant organisms, which has led to parallel reconstructions for *Saccharomyces cerevisiae*, *Yarrowia lipolytica*, *Scheffersomyces stipitis*, *Komagataella phaffii*, *Aspergillus niger* (Lopes and Rocha, 2017). Third, anchoring and availability biases cause non-conventional organisms’ GENRE’ s to be skewed towards model organism’ s metabolism while neglecting their own unique metabolism (Figure 1); examples of these commission and omission errors in yeast/fungi GENRE’ s can be found in the Supplementary Information. Finally, eukaryotic GENRE’ s have additional challenges not found in prokaryotes since they have larger genomes, few operons/gene clusters, numerous paralogs, alternative splicing, and compartments. These quantity and quality issues for GENRE hinders our ability to understand and control metabolism throughout the tree of life and highlight the need to build upon the bottom-up GENRE protocol outlined by Thiele and Palsson (2010).

**Figure 1.**
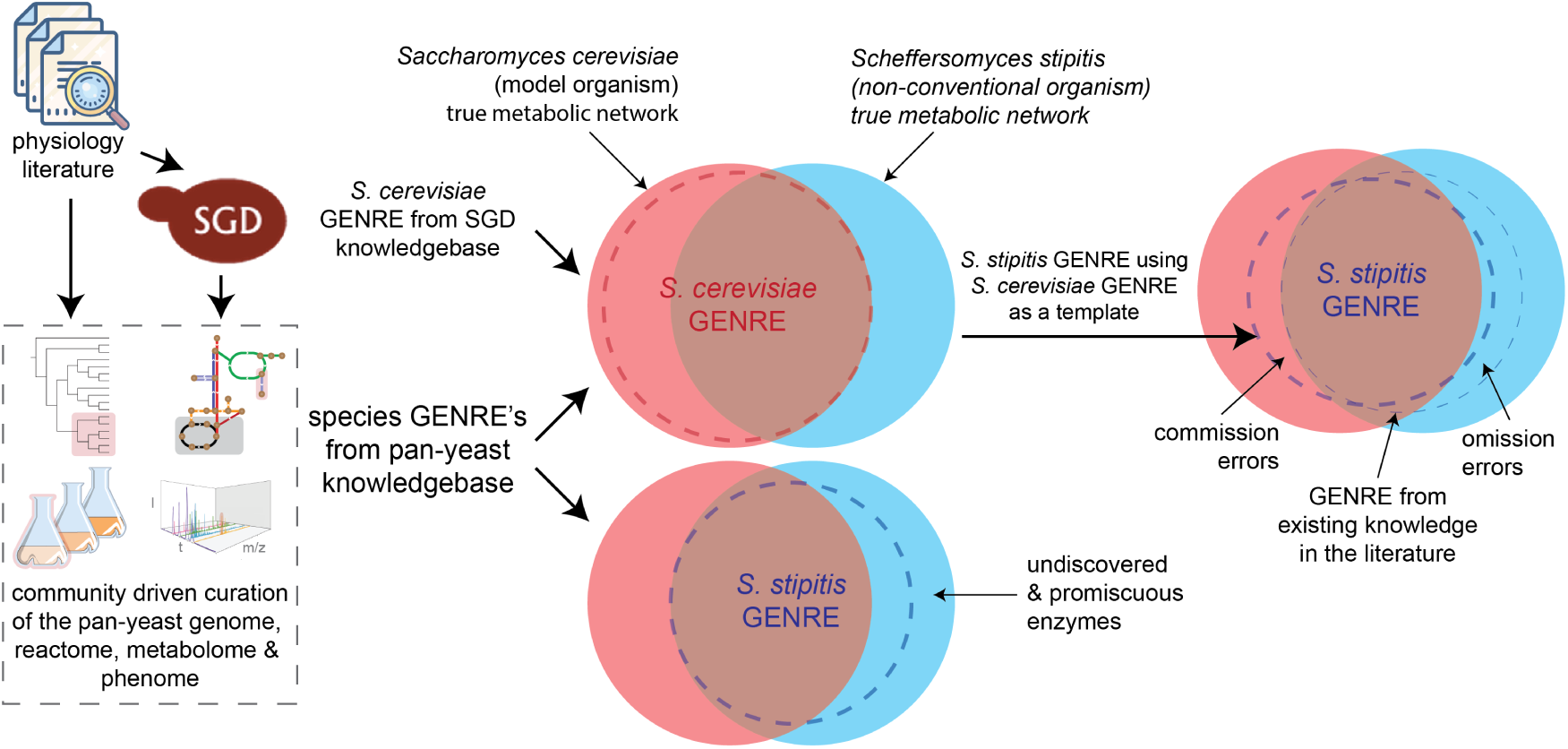
Non-conventional organism genome-scale network reconstruction (GENRE) using a model organism GENRE as a template versus databases from community-driven curation of a pan-genome, pan-reactome, pan-metabolome, and pan-phenome. *Saccharomyces* Genome Database (SGD) mines the literature to describe the physiology of *Saccharomyces cerevisiae* (a model organism). This knowledgebase is leveraged to compile a *S*. *cerevisiae* GENRE, which is its state of the art metabolic network; the gap between its true metabolic network and the state of the art GENRE represents undiscovered and promiscuous enzymes. In the absence of a curated genome database for *Scheffersomyces stipitis* (a non-conventional organism), the *S*. *cerevisiae* GENRE is used as a template to guide the *S*. *stipitis* GENRE. Anchoring bias makes the *S*. *stipitis* GENRE skew towards the *S*. *cereivisae* GENRE with commission errors; enzymes that have been characterized in *S*. *cerevisiae*, such as cytosolic NADP-dependent acetaldehyde dehydrogenase, have been included in past *S*. *stipitis* reconstructions despite no evidence based on orthology or enzymology (Correia et al., 2017). Availability bias prevents the GENRE curators from describing metabolism that is unique to *S*. *stipitis* and not well studied or documented, leading to omission errors; *S*. *stipitis* shares alkane hydroxylase orthologs with *Candida tropicalis*, a known alkane degrader (Lebeault et al., 1971), and yet alkane degradation has never been described in any *S*. *stipitis* or yeast GENRE. Community-driven curation of a pan-genome, pan-reactome, pan-metabolome, and pan-phenome reduces anchoring and availability biases by removing *S*. *cerevisiae* as the focal point for comparative metabolic network reconstruction; orthology assignment from rigorous phylogenic analysis of gene families rather than error-prone error prone methods; capturing non-canonical reactions catalyzed by promiscuous enzymes.

GENRE curators must make hundreds or thousands of *ad hoc* decisions, often without traceability (Ravikrishnan and Raman, 2015), because existing databases for orthologs, reactions, metabolites, and phenotypes are not always accurate or complete. This approach prevents GENRE’ s from being scaled throughout the tree of life. To solve these problems, we outline our approach for pan-GENRE with the formalized curation of orthologs, reactions, metabolites, and phenotypes for a larger group of organisms by research communities (Figure 2A). Previous studies have used this pan-genome approach for different strains of the same species to reconstruct metabolic networks (Islam et al., 2010; Monk et al., 2013), but the pan-genome can be expanded to a broader range of organisms (Vernikos et al., 2015). The first step in our framework is the curation of orthologs, paralogs, ohnologs, and xenologs within a pan-genome; many ortholog databases are available but our analysis has shown their ortholog groups sometimes contain paralogs (Correia et al., 2017). The second step is compiling the canonical *and* non-canonical reactions within a taxon and annotating them with ortholog-protein-reaction associations (OPR’ s). This reaction database can be collected from various sources (Figure 2B); existing reaction databases generally focus on canonical metabolism and underrepresent the metabolic potential of enzymes (Notebaart et al., 2014). Third, curating the presence and dynamics of metabolites from a larger group of organisms can help lead to the discovery of new pathways and enzymes (Caudy et al., 2018), especially when the metabolites are not found within the S-matrix. One notable example is the discovery of riboneogenesis, which was detected by sedoheptulose-1,7-bisphosphate changes in a shb17 knockout strain (Clasquin et al., 2011). Finally, curation of phenotypic data into machine-readable formats can help test, validate and improve GENRE’ s; Experiment Data Depot (EDD) has been recently described (Morrell et al., 2017) but this area is less developed than other databases. The primary purpose of these databases is for generating high-quality GENRE’ s, but they can also benefit the broader scientific community.

**Figure 2.**
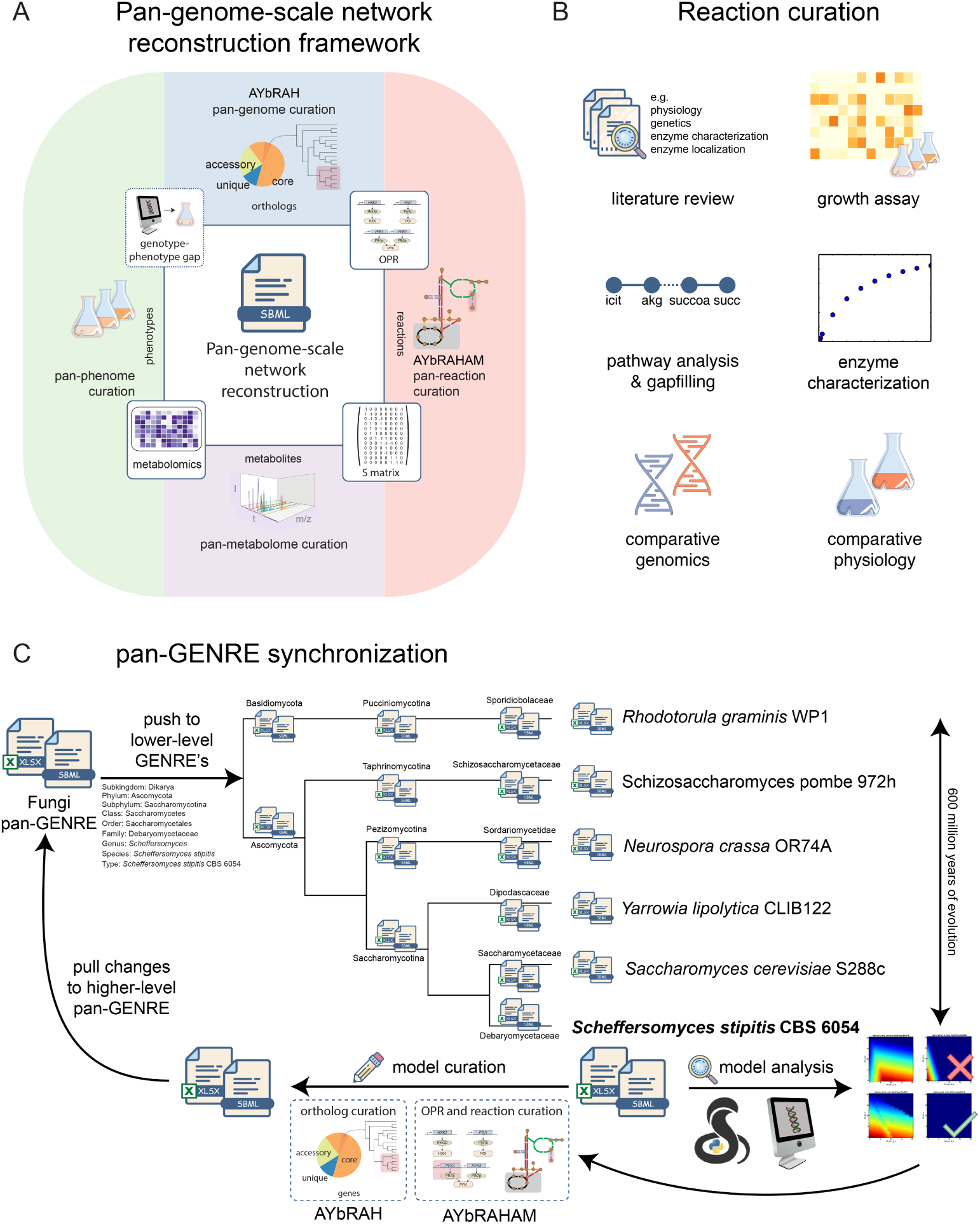
Pan-genome-scale network reconstruction framework for improving the quantity and quality of GENRE’ s. (A) A research community for a taxon, such as yeasts and fungi, curate its pan-genome, pan-reactome, pan-metabolome, and pan-phenome. Orthologs, paralogs, ohnologs, and xenologs are identified in the pan-genome by rigorous phylogenetic analysis. Reactions catalyzed by enzymes within a taxon are described in the pan-reactome and annotated with ortholog-protein-reaction associations (OPR). The presence and dynamics of metabolites are described in the pan-metabolome of a taxon; metabolites in the pan-metabolome not captured in the S-matrix can lead to new pathway discovery in the pan-reactome. Phenotypes within the taxon are transcribed into machine-readable formats to test, validate and improve genome-scale metabolic models. These databases are the basis for the pan-GENRE. (B) The pan-reactome is created by mining the literature, inferred reactions from growth assays and Biolog, pathway analysis and gap-filling, *in vitro* enzyme characterization and crude enzyme assays, comparative genomics and comparative physiology. (C) Our Fungi pan-GENRE spans 600 million years of evolution for 33 yeasts and fungi. It was used as a template to compile GENRE’ s for each taxonomic rank from sub-kingdom to strain taxonomic ranks. GENRE’ s for any taxonomic rank can by curated by updating the AYbRAH ortholog and AYbRAHAM reaction databases. These changes are pulled to the Fungi pan-GENRE, and pushed to the lower taxonomic rank GENRE’ s, enabling all the GENRE’ s to be synchronized. A GENRE can be used for genome-scale metabolic modelling (GSMM) to test new pathways and reconcile *in silico* and *in vivo* data. Further GENRE curation can bridge the gap between *in silico* and *in vivo* data.

We demonstrate our framework with a pan-GENRE for 33 yeasts/fungi spanning more than 600 million years of evolution. It was created by integrating AYbRAH, our yeast/fungal ortholog database, and Analyzing Yeasts by Reconstructing Ancestry of Homologs about Metabolism (AYbRAHAM), our new yeast/fungal reaction database. Our approach enables GENRE’ s for lower taxonomic levels, such as phylum, family or stains, to be compiled from the fungal pan-GENRE, and permits GENRE’ s to be synchronized with each other as the community characterizes more enzymes or research teams improve individual strain GENRE’ s (Figure 2C). We discuss the benefits and drawbacks of our approach and compare it with Model SEED (Devoid et al., 2013), CoReCo (Pitkänen et al., 2014), and CarveMe (Machado et al., 2018), existing methods for automated metabolic network reconstruction.

### 2 METHODS

A draft consensus pan-GENRE for Fungi was compiled from the following GENRE’ s using BiGG nomenclature (King et al., 2015): *Schizosaccharomyces pombe* (Sohn et al., 2012), *A*. *niger* (Andersen et al., 2008), *Y*. *lipolytica* (Pan and Hua, 2012; Loira et al., 2012), *K*. *phaffii* (Caspeta et al., 2012), *S*. *stipitis* (Balagurunathan et al., 2012; Caspeta et al., 2012; Li, 2012; Liu et al., 2012), *Eremothecium gossypii* (Ledesma-Amaro et al., 2014), and *S*. *cerevisiae* (Aung et al., 2013). Additional reactions were added to the pan-GENRE by reviewing enzyme assays, and reactions inferred from growth assays and Biolog data; these are referenced in the notes and reference section of the GENRE. Reactions were annotated with ortholog-protein-reaction associations (OPR’ s) using fungal ortholog groups (FOG’ s) from AYbRAH (Correia et al., 2017) if there was strong genomic evidence. OPR’ s for protein complexes were reviewed manually to determine the essential subunits. Protein complexes for *S*. *cerevisiae* and *S*. *pombe* were cross-referenced with the Complex Portal (Meldal et al., 2014). Reactions were also annotated with ORPHAN FOG if the reaction is catalyzed by an orphan enzyme and has a prerequisite FOG; ORPHAN RXN if the reaction has a prerequisite reaction; an organism code if there is evidence in a species or strain but the enzyme is unknown. Reactions annotated with BIOMASS, ESSENTIAL, ORPHAN, SPONTANEOUS, EQUILIBRIUM, DIFFUSION, EXCHANGE, or DEMAND were automatically included in all children GENRE’ s. A Python script was used to compile GENRE’ s for taxonomic ranks below Fungi in SBML3 and XLSX using the reaction annotations. We identified reactions requiring charge/mass balancing in the Fungi pan-GENRE with Memote (Lieven et al., 2018). PaperBLAST was used to find evidence of reactions for reactions in some ortholog groups (Price and Arkin, 2017).

### Biomass equations

The biomass equation was composed of protein, RNA, DNA, phospholipids, sterols, chitin, β -D-glucan, glycogen, trehalose, mannose. Each macromolecule was converted to a new metabolite species having a molar mass of 1000 g *ŀ* mmol^-1^ (see Supplementary Information). These macromolecule species enable the coefficients in the biomass equation to be set to the weight fraction of each macromolecule as measured in experiments; a similar approach was previously employed with SpoMBEL1693 (Sohn et al., 2012). The DNA composition for each strain GENRE was adjusted according to its GC content. Two protein synthesis reactions were constructed in the pan-GENRE: condensation from amino acids without a GTP cost and condensation of amino acids from charged tRNAs and with an additional cost of 2 mol GTP per mol amino acid (Schimmel, 1993). FOG’ s for the ribosome, RNA polymerase, DNA polymerase were assigned to the protein, RNA, and DNA biosynthesis reactions, respectively.

### FBA simulations

Flux balance analysis (FBA) was carried out with COBRApy v0.9.1. Reactions enabling artificial proton motive force cycles with glucose as a carbon source were blocked in the pan-GENRE (Pereira et al., 2016). Glucose and ammonium were used as a carbon source and nitrogen source to determine the yields of biomass precursors and correct any auxotrophies in all the strain GENRE’ s. Evolview v2 was used to map GENRE statistics and amino acid yields onto the species tree (He et al., 2016).

## 3 RESULTS

The Fungi pan-GENRE contains 1547 orthologs, 2726 reactions, 2226 metabolites, and 10 compartments. GENRE statistics for each strain are shown in Figure 3. Genes for the strain GENRE’ s range from 942 in *H*. *valbyensis* to 1558 in *Aspergillus niger*. The top subsystem in the strain GENRE’ s are biomass, transporters, amino acid metabolism, and central metabolism. Reaction and gene counts by subsystem for each strain GENRE is available in Figures S1 and S2.

**Figure 3.**
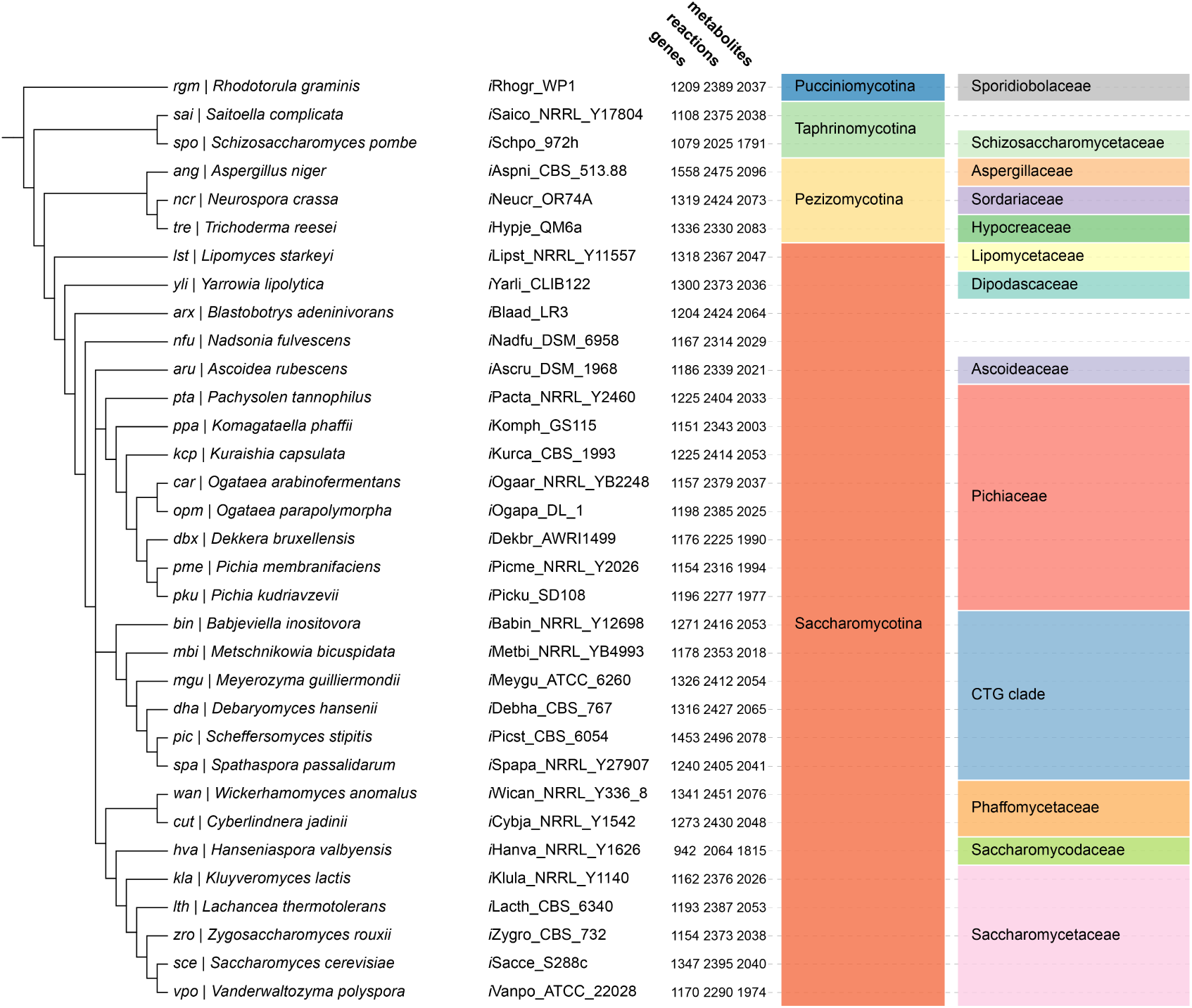
Yeast and fungi GENRE statistics. GENRE names, genes, reactions, metabolites, subphylum, and family mapped to a yeast/fungi species tree.

**Figure 4.**
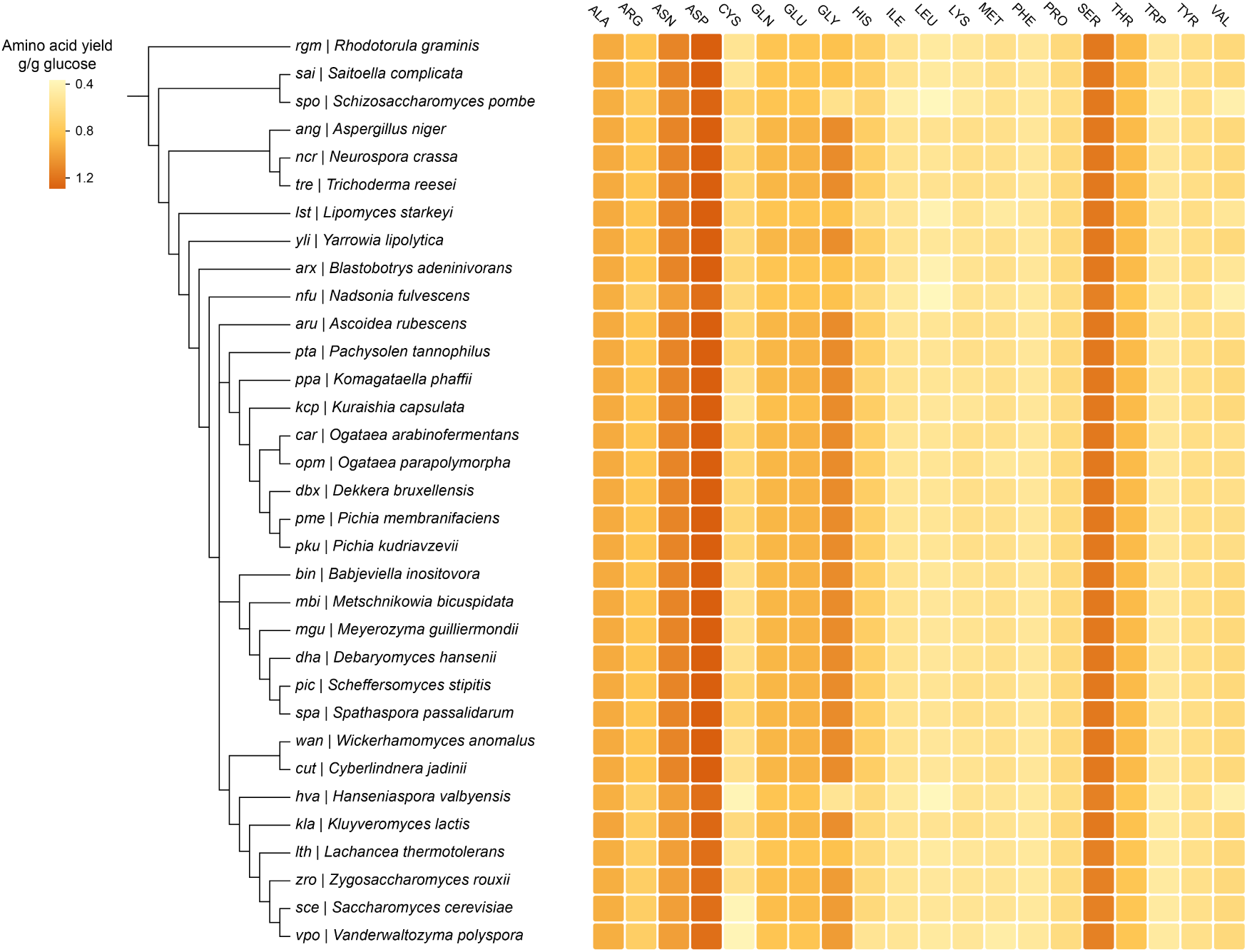
Heat map showing the yields of amino acids with glucose and ammonium as substrates with each strain genome-scale network reconstruction (GENRE). *Schizosaccharomyces pombe*, *Lipomyces starkeyi*, *Nadsonia fulvescens*, *Hanseniaspora valbyensis* all have reduced yields of glutamate, glutamine, glycine, histidine, leucine, methionine, and valine than the rest of the fungi and yeasts. *L*. *starkeyi* shares many genomic features as Proto-Yeast, the first budding yeast (Correia et al., 2017), while the remaining yeasts independently evolved the Crabtree effect. *S*. *pombe* and *H*. *valbyensis* have reduced genomes and primarily rely on glucose fermentation to ethanol.

### Expanded genomic coverage in the fungal pan-GENRE

Our strain GENRE’ s have more genes than GENRE’ s derived from manual curation or CoReCo (Table 1). Only yeastGEM and *i*MT1026, the most recent GENRE’ s for *S*. *cerevisiae* and *K*. *phaffii*, have more genes than our GENRE’ s when we compared the reconstructions without genes from the ribosome, RNA polymerase, DNA polymerase genes. These genes are not usually included in GENRE’ s. A quick inspection of the genes in *i*MT1026 indicates that it also includes RNA polymerase genes and uncharacterized genes/enzymes. Curating ortholog groups and annotating reactions with OPR’ s minimizes the chance that distantly related homologous genes are not added to GENRE’ s, and orthologous genes with low sequence similarity are captured in GENRE’ s.

**Table 1.**
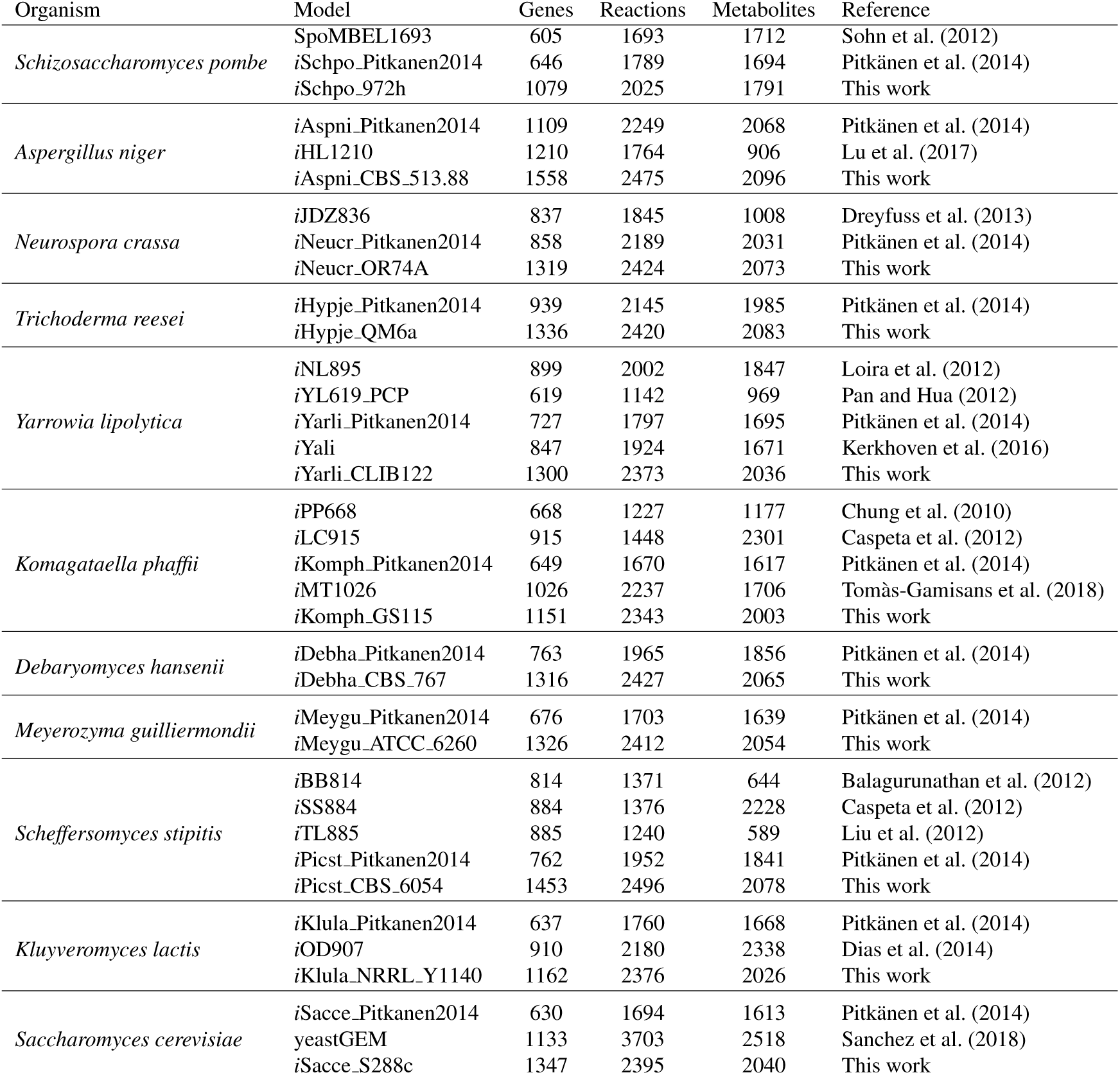
Comparison of genes, reactions, and metabolites in yeast and fungal GENRE’ s. AYbRAHAM GENRE’ s have more genes and reactions than GENRE’ s from manual curation or automatic reconstruction, with the exception of the reaction count in *S*. *cerevisiase*.

| Organism | Model | Genes | Reactions | Metabolites | Reference |
| --- | --- | --- | --- | --- | --- |
| *Schizosaccharomyces pombe* | SpoMBEL1693 | 605 | 1693 | 1712 | Sohn et al. (2012) |
|  | iSchpo_Pitkanen2014 | 646 | 1789 | 1694 | Pitkänen et al. (2014) |
|  | iSchpo_972h | 1079 | 2025 | 1791 | This work |
| *Aspergillus niger* | iAspni_Pitkanen2014 | 1109 | 2249 | 2068 | Pitkänen et al. (2014) |
|  | iHL1210 | 1210 | 1764 | 906 | Lu et al. (2017) |
|  | iAspni_CBS_513.88 | 1558 | 2475 | 2096 | This work |
| *Neurospora crassa* | iJDZ836 | 837 | 1845 | 1008 | Dreyfuss et al. (2013) |
|  | iNeucr_Pitkanen2014 | 858 | 2189 | 2031 | Pitkänen et al. (2014) |
|  | iNeucr_OR74A | 1319 | 2424 | 2073 | This work |
| *Trichoderma reesei* | iHypje_Pitkanen2014 | 939 | 2145 | 1985 | Pitkänen et al. (2014) |
|  | iHypje_QM6a | 1336 | 2420 | 2083 | This work |
| *Yarrowia lipolytica* | iNL895 | 899 | 2002 | 1847 | Loira et al. (2012) |
|  | iYL619_PCP | 619 | 1142 | 969 | Pan and Hua (2012) |
|  | iYarli_Pitkanen2014 | 727 | 1797 | 1695 | Pitkänen et al. (2014) |
|  | iYali | 847 | 1924 | 1671 | Kerkhoven et al. (2016) |
|  | iYarli_CLIB122 | 1300 | 2373 | 2036 | This work |
| *Komagataella phaffii* | iPP668 | 668 | 1227 | 1177 | Chung et al. (2010) |
|  | iLC915 | 915 | 1448 | 2301 | Caspeta et al. (2012) |
|  | iKomph_Pitkanen2014 | 649 | 1670 | 1617 | Pitkänen et al. (2014) |
|  | iMT1026 | 1026 | 2237 | 1706 | Tomàs-Gamisans et al. (2018) |
|  | iKomph_GS115 | 1151 | 2343 | 2003 | This work |
| *Debaryomyces hansenii* | iDebha_Pitkanen2014 | 763 | 1965 | 1856 | Pitkänen et al. (2014) |
|  | iDebha_CBS_767 | 1316 | 2427 | 2065 | This work |
| *Meyerozyma guilliermondii* | iMeygu_Pitkanen2014 | 676 | 1703 | 1639 | Pitkänen et al. (2014) |
|  | iMeygu_ATCC_6260 | 1326 | 2412 | 2054 | This work |
| *Scheffersomyces stipitis* | iBB814 | 814 | 1371 | 644 | Balagurunathan et al. (2012) |
|  | iSS884 | 884 | 1376 | 2228 | Caspeta et al. (2012) |
|  | iTL885 | 885 | 1240 | 589 | Liu et al. (2012) |
|  | iPicst_Pitkanen2014 | 762 | 1952 | 1841 | Pitkänen et al. (2014) |
|  | iPicst_CBS_6054 | 1453 | 2496 | 2078 | This work |
| *Kluyveromyces lactis* | iKlula_Pitkanen2014 | 637 | 1760 | 1668 | Pitkänen et al. (2014) |
|  | iOD907 | 910 | 2180 | 2338 | Dias et al. (2014) |
|  | iKlula_NRRL_Y1140 | 1162 | 2376 | 2026 | This work |
| *Saccharomyces cerevisiae* | iSacce_Pitkanen2014 | 630 | 1694 | 1613 | Pitkänen et al. (2014) |
|  | yeastGEM | 1133 | 3703 | 2518 | Sanchez et al. (2018) |
|  | iSacce_S288c | 1347 | 2395 | 2040 | This work |

### Expanded metabolic coverage in the fungal pan-GENRE

Our pan-GENRE has a wider metabolic coverage than prior yeast and fungal GENRE’ s. The pan-GENRE includes the degradation of alkanes and aromatics, which have been studied in Fungi for decades (Middelhoven et al., 1991, 1992; Van Beilen and Funhoff, 2007), and the degradation of aliphatic amines, which has been recently studied in *S*. *stipitis* by Linder (2014, 2018). These pathways are mediated by hydroxylases, monooxygenases, and dioxygenases from the cytochrome P450’ s family, which do not have well described gene-protein-reaction associations in current reaction databases. Enzymes in the cytochrome P450 family are known to be promiscuous and likely have vast unexplored metabolic capabilities in yeasts and fungi. Non-canonical metabolic reactions were also added to the GENRE, such as reactions catalyzed by ene-reductase. These reactions have no obvious role in metabolism (Zhang et al., 2016), but can be useful to know when engineering pathways in microbial hosts. The expanded metabolic coverage in our yeast and fungal GENRE’ s highlight the benefit for curated reaction and ortholog databases.

### Amino acid yields vary across yeast strains

The amino acid yields from glucose and ammonium vary in yeasts. There are strikingly lower yields in *S*. *pombe*, *Lipomyces starkeyi*, *Nadsonia fulvescens*, and *Hanseniaspora valbyensis* for glutamate, glutamine, glycine, histidine, leucine, methionine, and valine. *L*. *starkeyi* has many genomic and phenotypic features shared with Proto-Yeast, the first budding yeast (Correia et al., 2017), while the remaining yeasts independently evolved the Crabtree-effect (Correia et al., 2017). The overall protein yield, assuming the amino acid composition of *S*. *cerevisiae* from *i*MM904 (Mo et al., 2009), is lowest in yeasts that lost Complex I. The decrease in protein yield for these yeasts indicate that microbes are not always maximizing growth rate or biomass yield, and additional constraints (Bachmann et al., 2017) or environmental conditions can lead to loss of function. The differences in yield highlight the need to select microbial hosts with the highest potential for producing target biochemicals in metabolic engineering, or engineering higher yielding pathways in preferred hosts (Meadows et al., 2016). These hosts or pathways can be identified by running metabolic simulations with GENRE’ s from higher taxonomic ranks. Reconstructing GENRE’ s from curated databases can avoid commission errors that would lead to yields converging to the yields found in *S*. *cerevisiae* or other model organisms.

## 4 DISCUSSION

Most reaction databases reflect canonical metabolism, which understates the true metabolic capability of enzymes. AYbRAHAM includes non-canonical reactions identified in enzyme assays, which are relevant to metabolic engineering and enzyme evolution. Engineering the phosphoketolase pathway in *S*. *cerevisiae* required the deletion of *GPP1* and *GPP2*, which encode glycerol 3-phosphatase phosphatase, but have promiscuous acetyl-phosphate phosphatase activity (Hawkins et al., 2016). Acetyl-phosphate phosphatase would be considered a blocked reaction in a *S*. *cerevisiae* genome-scale metabolic model, even though this enzyme activity is important to metabolic engineers. Another example can be seen with rescuing a serine auxotrophy in *E*. *coli* from promiscuous phosphatase (Yip and Matsumura, 2013). Although these promiscuous activities are important to biologists and metabolic engineers, non-canonical reactions can increase the computational burden in genome-scale metabolic modelling. Reactions will be identified as core canonical to enable core metabolic network reconstructions to be compiled for every strain in future versions of the pan-GENRE. These examples highlight the need to capture the true metabolic network of microbes beyond canonical metabolism.

GENRE involves a review of the literature to capture metabolism unique to strains (österlund et al., 2012). One limitation of this approach is that a literature review based on *S*. *stipitis* would ignore metabolism it shares with other non-conventional organisms but has not been characterized directly, such as alkane degradation. A high-quality ortholog and reaction database with OPR’ s with our framework enables discoveries in one organism to be pushed to the GENRE’ s of all organisms within a taxon (Figure 2C). This synchronization can reduce the parallel GENRE efforts that have hampered the yeast community (Lopes and Rocha, 2017).

Our pan-GENRE framework relies on the ortholog conjecture, which states that orthologs are likely to have conserved function (Koonin, 2005); however, biological function is not always conserved amongst orthologs. For example, *NDE1* encodes external alternative NADH dehydrogenase and is present in all yeasts 33 yeasts and fungi in this study (Correia et al., 2017). Nde1p has been found to oxidize NADH and NADPH in several species (Bruinenberg et al., 1985; Melo et al., 2001; Overkamp et al., 2002; Miranda, 2011), but this ability was lost in the ancestor of *S*. *cerevisiae*. Furthermore, *S*. *pombe* appears to encode internal and external NADH dehydrogenase from the same gene (Crichton et al., 2007; Correia et al., 2017). To resolve these issues of non-conserved function in ortholog groups, we created new “functional ortholog groups” to assign these exceptions. Future methods can address these issues by assigning biological function to internal nodes in phylogenetic trees, or annotating reactions with transcripts instead of genes (Pfau et al., 2015).

### Comparison of pan-GENRE framework vs automated reconstruction

Model SEED has is commonly used to automate or semi-automate metabolic network reconstruction (Magnúsdóttir et al., 2017). Our experience with automated reconstruction using Model SEED has found that these models require extensive curation to address commission and omission errors (Correia et al., 2018), especially with enzyme complexes which are not fully annotated by Model SEED. Stringent assignment of orthology from phylogenetic analysis within a defined taxonomic group ensures these errors are minimized.

CoReCo is the first method to consider the simultaneous reconstruction of GENRE’ s for a wide range of species (Pitkänen et al., 2014). It relies on a probabilistic model to identify orthologs using bidirectional best hits with BLAST, which can complicate or misidentify orthology relationships for gene families with many duplications or high sequence similarity (Dalquen and Dessimoz, 2013; Correia et al., 2017). CoReCo was better at capturing reactions that were omitted by manual curation in past reconstructions, such as ATP citrate lyase in *S*. *pombe* (Sohn et al., 2012). On average, GENRE’ s from our framework have 74% more genes than CoReCo GENRE’ s (Table 1). CoReCo is an important milestone in expanding GENRE to diverse species, but curated ortholog and reaction databases are needed to ensure they are complete and accurate.

CarveMe is a top-down approach recently outlined by Machado et al. (2018), which is similar to our framework. They introduce the concept of a universal model for bacteria which can build GENRE’ s by homology using DIAMOND (Buchfink et al., 2014), or orthology via eggNOG-mapper (Huerta-Cepas et al., 2017). Using the GENRE’ s from 23 species, they were able to scale their universal model to more than 5000 bacterial GENRE’ s. This method can lead to several complications with eukaryotes, which have more paralogs than prokaryotes (Makarova et al., 2005). For example, *Aspergillus nidulans* encodes mitochondrial, cytosolic, and peroxisomal isocitrate dehydrogenase from IdpA, but most budding yeasts have these enzymes encoded by two or three genes (Szewczyk et al., 2001; Correia et al., 2017). Assigning a metabolic function to IdpA via orthology from *S*. *cerevisiae* would annotate it is a mitochondrial NADP-dependent isocitrate dehydrogenase, and omit its cytosolic and peroxisomal localizations. Therefore, homolog and ortholog-based assignments can both lead to incorrect gene-reaction-associations (GPR’ s) and enzyme localization. Furthermore, errors in genome annotations and existing GENRE’ s can be inherited in GENRE’ s derived from CarveMe. Our analysis of *S*. *stipitis*’ electron transport chain indicates that its genome does not encode internal alternative NADH dehydrogenase (Correia et al., 2017); however, the genome annotation of *S*. *stiptis* and *i*BB814 includes internal NADH dehydrogenase, encoded by PICST 58800 (Jeffries et al., 2007; Balagurunathan et al., 2012). These cases support the need for community consensus on orthologs, reaction networks, and OPR’ s for a group of organisms, which can compile high-quality GENRE’ s for lower-taxonomic levels.

## 5 CONCLUSIONS

The heart of GENRE process involves integrating information about orthologs and reactions, and to a lesser extent metabolites and phenotypes. The lack of complete and accurate ortholog and reaction databases forces research teams to make *ad hoc* decisions about the presence or absence of reactions, especially in non-conventional organisms. This time-consuming process limits the quantity and quality of GENRE’ s. We introduce our pan-GENRE framework to resolve these issues and demonstrate it with the metabolic network reconstruction of 33 yeasts and fungi, spanning 600 million years of evolution.

These GENRE’ s have more genomic and metabolic coverage than previous yeast and fungi GENRE’ s. The unified ortholog and reaction nomenclature in our GENRE’ s enables them to be synchronized as the pan-GENRE, or strain GENRE’ s are improved (Figure 2C). This approach can be scaled to other groups of organisms throughout the tree of life to increase the quantity and quality of GENRE’ s.

## 6 DATA AVAILABILITY

The AYbRAH ortholog database is available at https://github.com/kcorreia/aybrah. The AYbRAHAM reaction database, the code used to compile the GENRE’ s, and all GENRE’ s for each taxonomic rank in Fungi are available at https://github.com/kcorreia/aybraham.

## 8 ABBREVIATIONS

AYbRAH: Analyzing Yeasts by Reconstructing Ancestry of Homologs
AYbRAHAM: Analyzing Yeasts by Reconstructing Ancestry of Homologs about Metabolism EDD Experimental Data Depot
FOG: fungal ortholog group
GENRE: genome-scale network reconstruction GPR gene-protein-reaction associations
OPR: ortholog-protein-reaction association
pan-GENRE: pan-genome-scale network reconstruction

## REFERENCES

Andersen, M. R., Nielsen, M. L., and Nielsen, J. (2008). Metabolic model integration of the bibliome, genome, metabolome and reactome of Aspergillus niger. Molecular Systems Biology, 4(1):178.

Aung, H. W., Henry, S. A., and Walker, L. P. (2013). Revising the representation of fatty acid, glycerolipid, and glycerophospholipid metabolism in the consensus model of yeast metabolism. Industrial Biotechnology, 9(4):215–228.

Bachmann, H., Molenaar, D., Branco dos Santos, F., and Teusink, B. (2017). Experimental evolution and the adjustment of metabolic strategies in lactic acid bacteria. FEMS Microbiology Reviews, page fux024.

Balagurunathan, B., Jonnalagadda, S., Tan, L., and Srinivasan, R. (2012). Reconstruction and analysis of a genome-scale metabolic model for Scheffersomyces stipitis. Microbial Cell Factories, 11(1):1.

Bruinenberg, P. M., van Dijken, J. P., Kuenen, J. G., and Scheffers, W. A. (1985). Oxidation of NADH and NADPH by mitochondria from the yeast Candida utilis. Microbiology, 131(5):1043–1051.

Buchfink, B., Xie, C., and Huson, D. H. (2014). Fast and sensitive protein alignment using DIAMOND. Nature Methods, 12(1):59.

Caspeta, L., Shoaie, S., Agren, R., Nookaew, I., and Nielsen, J. (2012). Genome-scale metabolic reconstructions of Pichia stipitis and Pichia pastoris and in silico evaluation of their potentials. BMC Systems Biology, 6(1):1.

Caudy, A. A., Hanchard, J. A., Hsieh, A., Shaan, S., and Rosebrock, A. P. (2018). Functional genetic discovery of enzymes using full-scan mass spectrometry metabolomics. Biochemistry and Cell Biology, (ja).

Chung, B. K., Selvarasu, S., Camattari, A., Ryu, J., Lee, H., Ahn, J., Lee, H., and Lee, D.-Y. (2010). Genome-scale metabolic reconstruction and in silico analysis of methylotrophic yeast Pichia pastoris for strain improvement. Microbial Cell Factories, 9(1):50.

Clasquin, M. F., Melamud, E., Singer, A., Gooding, J. R., Xu, X., Dong, A., Cui, H., Campagna, S. R., Savchenko, A., Yakunin, A. F., et al. (2011). Riboneogenesis in yeast. Cell, 145(6):969–980.

Correia, K., Ho, H., and Mahadevan, R. (2018). Genome-scale metabolic network reconstruction of the chloroform-respiring Dehalobacter restrictus strain CF. bioRxiv, page 375063.

Correia, K., Yu, S. M., and Mahadevan, R. (2017). Reconstructing the evolution of metabolism in budding yeasts. bioRxiv, page 237974.

Crichton, P. G., Affourtit, C., and Moore, A. L. (2007). Identification of a mitochondrial alcohol dehydrogenase in Schizosaccharomyces pombe: new insights into energy metabolism. Biochemical Journal, 401(2):459–464.

Dalquen, D. A. and Dessimoz, C. (2013). Bidirectional best hits miss many orthologs in duplication-rich clades such as plants and animals. Genome Biology and Evolution, 5(10):1800–1806.

Devoid, S., Overbeek, R., DeJongh, M., Vonstein, V., Best, A. A., and Henry, C. (2013). Automated genome annotation and metabolic model reconstruction in the SEED and Model SEED. In Systems Metabolic Engineering, pages 17–45. Springer.

Dias, O., Pereira, R., Gombert, A. K., Ferreira, E. C., and Rocha, I. (2014). iOD907, the first genome-scale metabolic model for the milk yeast Kluyveromyces lactis. Biotechnology Journal, 9(6):776–790.

Dreyfuss, J. M., Zucker, J. D., Hood, H. M., Ocasio, L. R., Sachs, M. S., and Galagan, J. E. (2013). Reconstruction and validation of a genome-scale metabolic model for the filamentous fungus Neurospora crassa using FARM. PLoS Computational Biology, 9(7):e1003126.

Hackett, S. R., Zanotelli, V. R., Xu, W., Goya, J., Park, J. O., Perlman, D. H., Gibney, P. A., Botstein, D., Storey, J. D., and Rabinowitz, J. D. (2016). Systems-level analysis of mechanisms regulating yeast metabolic flux. Science, 354(6311).

Hawkins, K. M., Mahatdejkul-Meadows, T. T., Meadows, A. L., Pickens, L. B., Tai, A., and Tsong, A. E. (2016). Use of phosphoketolase and phosphotransacetylase for production of acetyl-coenzyme A derived compounds. US Patent 9, 410, 214.

He, Z., Zhang, H., Gao, S., Lercher, M. J., Chen, W.-H., and Hu, S. (2016). Evolview v2: an online visualization and management tool for customized and annotated phylogenetic trees. Nucleic Acids Research, pages W236–241.

Huerta-Cepas, J., Forslund, K., Pedro Coelho, L., Szklarczyk, D., Juhl Jensen, L., von Mering, C., and Bork, P. (2017). Fast genome-wide functional annotation through orthology assignment by eggNOG-Mapper. Molecular Biology and Evolution, 34(8):2115–2122.

Islam, M. A., Edwards, E. A., and Mahadevan, R. (2010). Characterizing the metabolism of Dehalococcoides with a constraint-based model. PLoS Computational Biology, 6(8):e1000887.

Jeffries, T. W., Grigoriev, I. V., Grimwood, J., Laplaza, J. M., Aerts, A., Salamov, A., Schmutz, J., Lindquist, E., Dehal, P., Shapiro, H., et al. (2007). Genome sequence of the lignocellulose-bioconverting and xylose-fermenting yeast Pichia stipitis. Nature Biotechnology, 25(3):319–326.

Kerkhoven, E. J., Pomraning, K. R., Baker, S. E., and Nielsen, J. (2016). Regulation of amino-acid metabolism controls flux to lipid accumulation in Yarrowia lipolytica. NPJ Systems Biology and Applications, 2:16005.

King, Z. A., Lu, J., Dräger, A., Miller, P., Federowicz, S., Lerman, J. A., Ebrahim, A., Palsson, B. O., and Lewis, N. E. (2015). BiGG models: A platform for integrating, standardizing and sharing genome-scale models. Nucleic Acids Research, 44(D1):D515–D522.

Koonin, E. V. (2005). Orthologs, paralogs, and evolutionary genomics. Annual Review of Genetics, 39:309–338.

Lebeault, J.-M., Lode, E. T., and Coon, M. J. (1971). Fatty acid and hydrocarbon hydroxylation in yeast: role of cytochrome P-450 in Candida tropicalis. Biochemical and Biophysical Research Communications, 42(3):413–419.

Ledesma-Amaro, R., Kerkhoven, E. J., Revuelta, J. L., and Nielsen, J. (2014). Genome scale metabolic modeling of the riboflavin overproducer Ashbya gossypii. Biotechnology and Bioengineering, 111(6):1191–1199.

Li, P. Y. (2012). In silico metabolic network reconstruction of Scheffersomyces stipitis. Master’s thesis, University of Toronto.

Lieven, C., Beber, M. E., Olivier, B. G., Bergmann, F. T., Babaei, P., Bartell, J. A., Blank, L. M., Chauhan, S., Correia, K., Diener, C., et al. (2018). Memote: A community-driven effort towards a standardized genome-scale metabolic model test suite. bioRxiv, page 350991.

Linder, T. (2014). CMO1 encodes a putative choline monooxygenase and is required for the utilization of choline as the sole nitrogen source in the yeast Scheffersomyces stipitis (syn. Pichia stipitis). Microbiology, 160(5):929–940.

Linder, T. (2018). Genetic redundancy in the catabolism of methylated amines in the yeast Scheffersomyces stipitis. Antonie van Leeuwenhoek, 111(3):401–411.

Liu, T., Zou, W., Liu, L., and Chen, J. (2012). A constraint-based model of Scheffersomyces stipitis for improved ethanol production. Biotechnology for Biofuels, 5(1):72.

Loira, N., Dulermo, T., Nicaud, J.-M., and Sherman, D. J. (2012). A genome-scale metabolic model of the lipid-accumulating yeast Yarrowia lipolytica. BMC Systems Biology, 6(1):35.

Lopes, H. and Rocha, I. (2017). Genome-scale modeling of yeast: chronology, applications and critical perspectives. FEMS Yeast Research, 17(5).

Lu, H., Cao, W., Ouyang, L., Xia, J., Huang, M., Chu, J., Zhuang, Y., Zhang, S., and Noorman, H. (2017). Comprehensive reconstruction and in silico analysis of Aspergillus niger genome-scale metabolic network model that accounts for 1210 ORFs. Biotechnology and Bioengineering, 114(3):685–695.

Machado, D., Andrejev, S., Tramontano, M., and Patil, K. R. (2018). Fast automated reconstruction of genome-scale metabolic models for microbial species and communities. Nucleic Acids Research.

Magnúsdóttir, S., Heinken, A., Kutt, L., Ravcheev, D. A., Bauer, E., Noronha, A., Greenhalgh, K., Jäger, C., Baginska, J., Wilmes, P., et al. (2017). Generation of genome-scale metabolic reconstructions for 773 members of the human gut microbiota. Nature Biotechnology, 35(1):81.

Makarova, K. S., Wolf, Y. I., Mekhedov, S. L., Mirkin, B. G., and Koonin, E. V. (2005). Ancestral paralogs and pseudoparalogs and their role in the emergence of the eukaryotic cell. Nucleic Acids Research, 33(14):4626–4638.

Meadows, A. L., Hawkins, K. M., Tsegaye, Y., Antipov, E., Kim, Y., Raetz, L., Dahl, R. H., Tai, A., Mahatdejkul-Meadows, T., Xu, L., et al. (2016). Rewriting yeast central carbon metabolism for industrial isoprenoid production. Nature, 537(7622):694–697.

Meldal, B. H., Forner-Martinez, O., Costanzo, M. C., Dana, J., Demeter, J., Dumousseau, M., Dwight, S. S., Gaulton, A., Licata, L., Melidoni, A. N., et al. (2014). The complex portal-an encyclopaedia of macromolecular complexes. Nucleic Acids Research, 43(D1):D479–D484.

Melo, A. M., Duarte, M., Møller, I. M., Prokisch, H., Dolan, P. L., Pinto, L., Nelson, M. A., and Videira, A. (2001). The external calcium-dependent NADPH dehydrogenase from Neurospora crassa mitochondria. Journal of Biological Chemistry, 276(6):3947–3951.

Middelhoven, W. J., Coenen, A., Kraakman, B., and Gelpke, M. D. S. (1992). Degradation of some phenols and hydroxybenzoates by the imperfect ascomycetous yeasts Candida parapsilosis and Arxula adeninivorans: evidence for an operative gentisate pathway. Antonie van Leeuwenhoek, 62(3):181–187.

Middelhoven, W. J., de Jong, I. M., and de Winter, M. (1991). Arxula adeninivorans, a yeast assimilating many nitrogenous and aromatic compounds. Antonie van Leeuwenhoek, 59(2):129–137.

Miranda, A. R. d. (2011). Deleçaõ do gene PGI1 da levedura Pichia stiptis para aumentar o rendimento fermentativo a etanol.

Mo, M. L., Palsson, B. Ø., and HerrgÅrd, M. J. (2009). Connecting extracellular metabolomic measurements to intracellular flux states in yeast. BMC Systems Biology, 3(1):37.

Monk, J., Nogales, J., and Palsson, B. O. (2014). Optimizing genome-scale network reconstructions. Nature Biotechnology, 32(5):447.

Monk, J. M., Charusanti, P., Aziz, R. K., Lerman, J. A., Premyodhin, N., Orth, J. D., Feist, A. M., and Palsson, B. Ø. (2013). Genome-scale metabolic reconstructions of multiple Escherichia coli strains highlight strain-specific adaptations to nutritional environments. Proceedings of the National Academy of Sciences, 110(50):20338–20343.

Morrell, W. C., Birkel, G. W., Forrer, M., Lopez, T., Backman, T. W., Dussault, M., Petzold, C. J., Baidoo, E. E., Costello, Z., Ando, D., et al. (2017). The Experiment Data Depot: a web-based software tool for biological experimental data storage, sharing, and visualization. ACS Synthetic Biology, 6(12):2248–2259.

Notebaart, R. A., Szappanos, B., Kintses, B., Pál, F., Györkei, á., Bogos, B., Lázár, V., Spohn, R., Csörgo, B., Wagner, A., et al. (2014). Network-level architecture and the evolutionary potential of underground metabolism. Proceedings of the National Academy of Sciences, 111(32):11762–11767.

österlund, T., Nookaew, I., Bordel, S., and Nielsen, J. (2013). Mapping condition-dependent regulation of metabolism in yeast through genome-scale modeling. BMC Systems Biology, 7(1):1.

österlund, T., Nookaew, I., and Nielsen, J. (2012). Fifteen years of large scale metabolic modeling of yeast: developments and impacts. Biotechnology Advances, 30(5):979–988.

Overkamp, K. M., Bakker, B. M., Steensma, H., van Dijken, J. P., and Pronk, J. T. (2002). Two mechanisms for oxidation of cytosolic NADPH by Kluyveromyces lactis mitochondria. Yeast, 19(10):813–824.

Pan, P. and Hua, Q. (2012). Reconstruction and in silico analysis of metabolic network for an oleaginous yeast, Yarrowia lipolytic. PLoS One, 7(12):e51535.

Pereira, R., Nielsen, J., and Rocha, I. (2016). Improving the flux distributions simulated with genome-scale metabolic models of Saccharomyces cerevisiae. Metabolic Engineering Communications, 3:153–163.

Pfau, T., Pacheco, M. P., and Sauter, T. (2015). Towards improved genome-scale metabolic network reconstructions: unification, transcript specificity and beyond. Briefings in Bioinformatics, 17(6):1060–1069.

Pitkänen, E., Jouhten, P., Hou, J., Syed, M. F., Blomberg, P., Kludas, J., Oja, M., Holm, L., Penttilä, M., Rousu, J., et al. (2014). Comparative genome-scale reconstruction of gapless metabolic networks for present and ancestral species. PLoS Computational Biology, 10(2):e1003465.

Price, M. N. and Arkin, A. P. (2017). PaperBLAST: text mining papers for information about homologs. mSystems, 2(4):e00039–17.

Ravikrishnan, A. and Raman, K. (2015). Critical assessment of genome-scale metabolic networks: the need for a unified standard. Briefings in Bioinformatics, 16(6):1057–1068.

Sanchez, B. J., Li, F., Kerkhoven, E. J., and Nielsen, J. (2018). SLIMEr: probing flexibility of lipid metabolism in yeast with an improved constraint-based modeling framework. bioRxiv, page 324863.

Schimmel, P. (1993). GTP hydrolysis in protein synthesis: two for Tu? Science, 259(5099):1264–1266.

Sohn, S. B., Kim, T. Y., Lee, J. H., and Lee, S. Y. (2012). Genome-scale metabolic model of the fission yeast Schizosaccharomyces pombe and the reconciliation of in silico/in vivo mutant growth. BMC Systems Biology, 6(1):1.

Szewczyk, E., Andrianopoulos, A., Davis, M. A., and Hynes, M. J. (2001). A single gene produces mitochondrial, cytoplasmic, and peroxisomal NADP-dependent isocitrate dehydrogenase in Aspergillus nidulans. Journal of Biological Chemistry, 276(40):37722–37729.

Thiele, I., Clancy, C. M., Heinken, A., and Fleming, R. M. (2017). Quantitative systems pharmacology and the personalized drug–microbiota–diet axis. Current Opinion in Systems Biology, 4:43–52.

Thiele, I. and Palsson, B. Ø. (2010). A protocol for generating a high-quality genome-scale metabolic reconstruction. Nature Protocols, 5(1):93.

Tomàs-Gamisans, M., Ferrer, P., and Albiol, J. (2018). Fine-tuning the P. pastoris iMT1026 genome-scale metabolic model for improved prediction of growth on methanol or glycerol as sole carbon sources. Microbial Biotechnology, 11(1):224–237.

Van Beilen, J. B. and Funhoff, E. G. (2007). Alkane hydroxylases involved in microbial alkane degradation. Applied Microbiology and Biotechnology, 74(1):13–21.

Vernikos, G., Medini, D., Riley, D. R., and Tettelin, H. (2015). Ten years of pan-genome analyses. Current Opinion in Microbiology, 23:148–154.

Yip, S. H.-C. and Matsumura, I. (2013). Substrate ambiguous enzymes within the Escherichia coli proteome offer different evolutionary solutions to the same problem. Molecular Biology and Evolution, 30(9):2001–2012.

Zhang, B., Zheng, L., Lin, J., and Wei, D. (2016). Characterization of an ene-reductase from Meyerozyma guilliermondii for asymmetric bioreduction of a, ß -unsaturated compounds. Biotechnology Letters, 38(9):1527–1534.

